# Transformation of human chondrocytes with copper-containing metal-organic biohybrids (MOBs)

**DOI:** 10.1101/2024.01.12.575456

**Authors:** Tasneem Khasru, Katie McKenzie, Kyle Rugg, Shaylee Boudreaux, Kelly McMahen, Navya Uppu, Mark A. DeCoster

**Affiliations:** Louisiana Tech University, Cellular Neuroscience Lab, Department of Biomedical Engineering, Ruston, LA, USA 71272; Louisiana Tech University, Cellular Neuroscience Lab, Institute for Micromanufacturing, Ruston, LA, USA 71272

**Keywords:** chondrocytes, fibroblasts, copper, collagen, cell morphology

## Abstract

Copper is involved in the biosynthesis of collagen, however soluble copper salts dissipate quickly and copper nanoparticles are cytotoxic. Here we added a novel copper-containing nanomaterial (CuHARS) to assess human chondrocyte function in the presence of copper. Human dermal fibroblasts (HDFs) were also treated as a control. Chondrocyte response to CuHARS was assessed by chronic nanomaterial treatment (30 µg/ml) followed by digital microscopy and image analysis of cellular features compared to normal chondrocytes. Unexpectedly, chronic CuHARS treatment of human chondrocytes transformed cells over time to cells with extremely elongated and variegated processes and lower proliferation rates compared to normal chondrocytes. In these transformed cells, which we named 3G, shedding of fine processes was observed over time and collected supernatants demonstrated elevated collagen levels compared to normal cell culture media. In contrast to chondrocytes, HDFs treated with CuHARS showed attenuated changes in morphology, and notably retained a prominent ability for continued proliferation. These results demonstrate that a copper-containing biohybrid material (CuHARS) can stably transform human chondrocytes with highly altered morphology, lower proliferation rates, and altered membrane dynamics compared to normal chondrocytes. In contrast, human dermal fibroblasts demonstrated attenuated changes in morphology, and retained an enhanced ability for proliferation.

## Introduction

Metal-organic biohybrids (MOBs) are a family of materials that self-assemble at physiological temperatures when combining metals including copper, silver, and cobalt, and the biological component of the amino acid dimer cystine [1, 2]. The copper-containing MOB forms as a linear, high-aspect ratio structure which we have named CuHARS for copper-high aspect ratio structures [3]. CuHARS scale from the nano-micro in dimensions, and are completely degradable under physiological conditions [4]. Considering the importance of copper in collagen formation [5-7], we hypothesized that CuHARS when combined with chondrocytes, might reveal a transformative function. Indeed we found here that chronic treatment with CuHARS could transform them into slowly proliferating cells with highly extended and branched processes. Over time these processes were shed from the cell bodies and supernatants collected from these cultures indicated elevated levels of collagen compared to media alone. Control treatments were carried out with human dermal fibroblasts and demonstrated attenuated changes in morphology compared to chondrocytes, but with retention on high proliferation rate compared to the transformed (3G) cells.

## Materials and Methods

### CuHARS

Copper-containing high-aspect ratio structures (CuHARS) were prepared as previously described [3, 4] and suspended in 70% isopropyl alcohol in sterile polypropylene tubes until applied to cells.

### Cell culture

Human chondrocytes and human dermal fibroblasts were obtained from Cell Applications (San Diego, CA, USA) and grown in cell culture flasks until the third passage at which point, they were treated with 30 μg/ml of CuHARS.

### Microscopy

Images were taken on a Leica DMI 6000B inverted microscope with phase or brightfield optics as indicated, and powered by version 2.0 acquisition and software.

### Image Analysis

Brightfield images of CuHARS bound to cells and degrading over time were post-processed and analyzed using Image Pro Plus software (Ver. 7.0) from Media Cybernetics (Silver Spring, MD, USA). The densely dark and elongated CuHARS provided high contrast for imaging and quantifying material associated with cells. The dark material was then used to create an inverted mask of white material on a black background, and these mask images used to quantify the change in aspect ratio of regions of interest over time (See **Figure 1** and **Supplementary Figure 1**).

**Figure 1.**
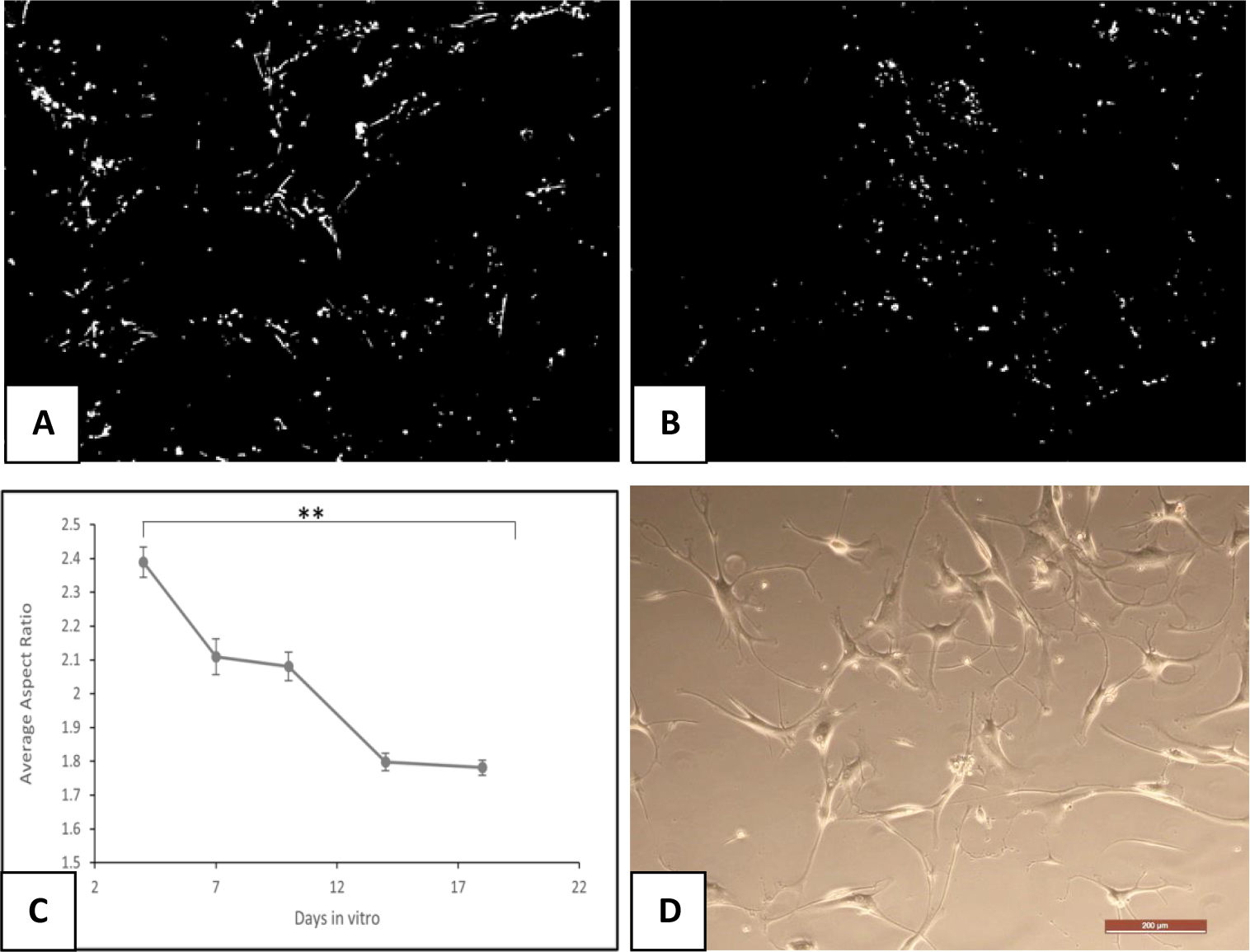
Masks of CuHARS associated with chondrocytes were generated as described in Methods at 4 DIV (A) and 18 DIV (B) using brightfield microscopy. The aspect ratio dynamics of CuHARS treatment on chondrocytes was analyzed (C). Data at each time point represent over 1,000 objects analyzed, with averages and standard error bars shown; P values (**)<0.01. (D) Transformed chondrocytes demonstrated variegated and elongated processes. Scale bar in D represents 200 microns, and masks generated in A and B are of the same magnification.

### Collagen Assay

A collagen assay kit was obtained from Sigma-Aldrich (cat# MAK322) to determine the amount of collagen in cell culture supernatants. Sample preparation, master mix preparation, and assay reactions were done according to instructions provided by the manufacturer with the following modifications: A total volume of 598 μl was used for each reading utilizing Perfector Scientific micro-cuvettes (Catalogue #9008). A wavelength scan was then carried out with excitation at 375 nm and emission from 465-499 nm with a peak detected at 465 nm using a Hitachi F-2500 fluorescence spectrophotometer. A sample of cell culture basal media was tested to determine the baseline fluorescence readings and then compared to samples collected from cellular supernatants and to pure collagen controls as indicated.

### Cryopreservation

Normal human chondrocytes and 3G transformed chondrocytes were cryopreserved in complete cell culture media with the addition of 10% fetal bovine serum (FBS) and 10% dimethyl sulfoxide (DMSO) as recommended by the vendor. Cells were then slowly frozen at -80° C in a isopropyl freezing jar overnight, followed by immersion in liquid nitrogen dewars.

## Results

To address the question of whether the novel copper-containing nanomaterial CuHARS could alter human chondrocyte function, we chronically treated flasks of cells with 30 µg/ml of CuHARS, tracking the materials for degradation and materials changes using digital microscopy and image analysis (**Figure 1A**). We had previously shown that CuHARS could be completely degraded under physiological conditions [4] in cell culture media only. Here we found that over 18 days in vitro (DIV), CuHARS materials associated with chondrocytes (**Supplementary Figure S1**), and significantly changed size and shape, shifting from elongated at the beginning to more particulate over time (**Figures 1A&B**). This system thus provided a label-free method for tracking cell changes. This combination of change in shape of the CuHARS (aspect ratio), and change in size over time, allowed for the generation of images of sorted identified objects (**Supplementary Figure S2**) created using Image Pro Plus software, which were used to track dynamic changes in the treated chondrocytes over 18 DIV (**Figure 1C**).

Upon collecting cells floating cells from the treated flasks and replating them, we discovered that the CuHARS treatment had transformed the cells as expressed by novel morphology, including extremely long and variegated cellular processes compared to normal human chondrocytes (**Figure 1D**). We named these transformed cells 3G cells.

Transformed 3G cells over time in culture shed membrane material from the elongated fine, and variegated processes (**Figure 2A**). In some cases, these transformed cells grew to be individually very large with prominent membrane shedding (**Figure 2B**).

**Figure 2.**
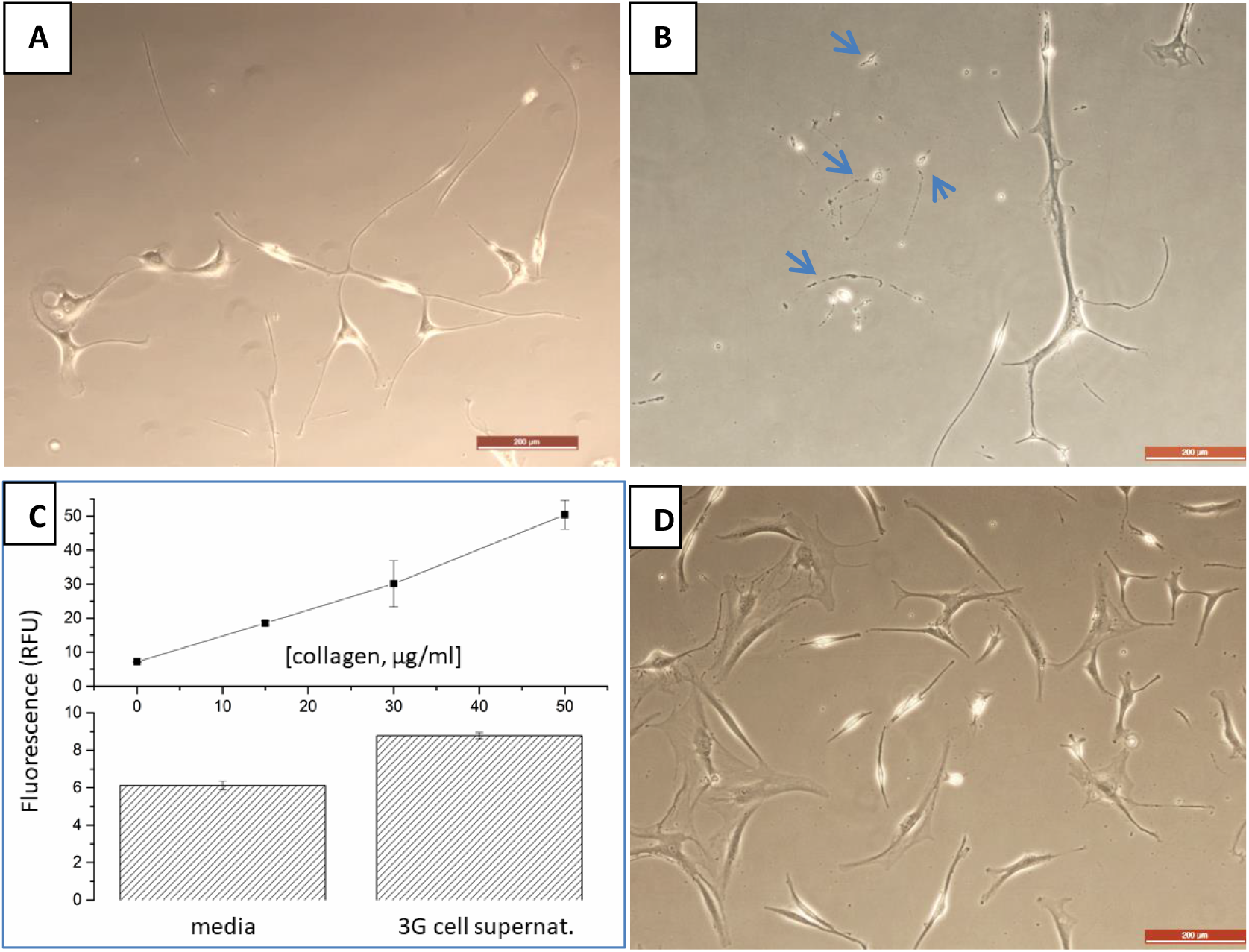
Cell membrane fragmentation and shedding with collagen formation. Transformed, 3G chondrocytes (A), demonstrate elongated processes and early breakage and shedding of processes. In some areas, transformed 3G cells were huge, spanning many hundreds of microns, with fine processes that showed numerous fragmentation areas (B); arrows indicate fragmented cell areas. Collected cell culture supernatants from 3G cells had elevated collagen levels compared to media alone as normalized to pure collagen controls (C). In contrast to 3G cells, normal chondrocytes had smaller processes, which remained contiguous with cell bodies and cells proliferated, forming dense areas over time (D). Scale bars indicate 200 microns in all panels; all cells shown at 12 DIV.

When supernatants were collected from 3G cultures and assayed for collagen, the levels for collagen measured were significantly elevated (1.4-fold increase) compared to culture media alone (**Figure 2C**). In contrast to the large cell size, extended processes, and membrane shedding of transformed 3G cells, normal chondrocytes remained smaller in size, and proliferated over time, demonstrating dense areas of many cells per field (**Figure 2D**). The highly elongated, and highly branched fine processes observed in transformed 3G cells were not observed in normal chondrocytes (**Figure S3**). Furthermore, both transformed 3G cells and normal chondrocytes could be maintained for extensive periods in vitro (40 DIV), with mature 3G cultures establishing extreme, highly branched processes with few cell bodies, while normal chondrocytes became highly dense with many cells, forming monolayers with few processes per cell (**Figure S4**).

To test for stable transformation of 3G cells, these cells were recombined with normal chondrocytes at low densities and dynamic growth tracked using digital microscopy. Distinct morphology of 3G cells remained apparent even when mixed with normal chondrocytes, with 3G cells maintaining large cell areas with fine extended processes compared to surrounding normal chondrocytes (**Figure 3**), and in some cases, breakage and fragmentation of fine processes was observed (Figure 3B), as was seen in 3G cell cultures alone (**Figure 2A**).

**Figure 3.**
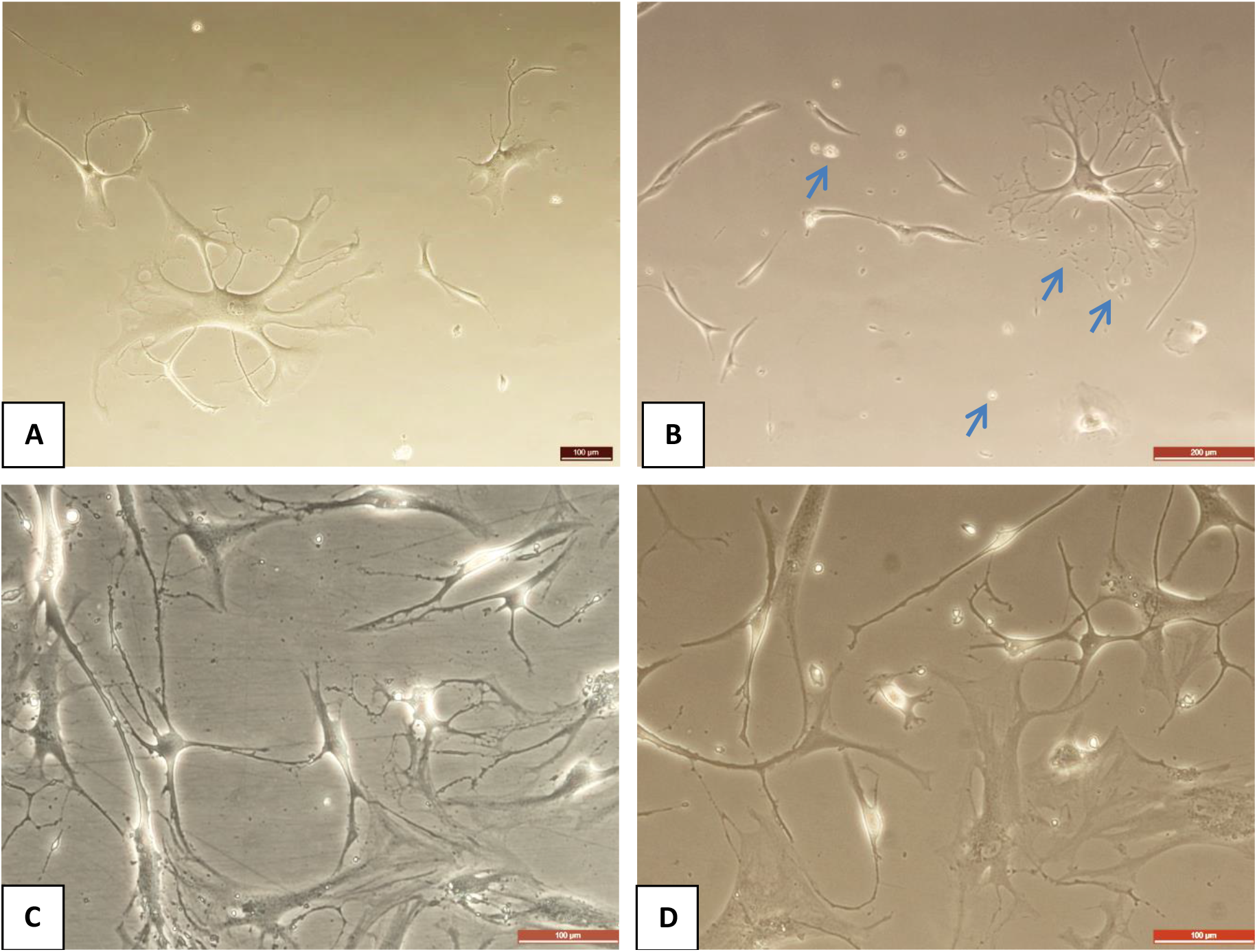
Combination of transformed 3G chondrocytes and normal chondrocytes in mixed cultures. Mixed cultures were documented at 3DIV (A, top row) and 14DIV (B, bottom row), respectively. Arrows in 3B indicated cell fragmentation areas. Scale bars indicated are 100 microns (A, C, and D) or 200 microns (B).

## Discussion

These results provide a method for transforming human chondrocytes with chronic treatment with copper in the form of the metal-organic biohybrid CuHARS. The results are novel, and unexpected, due to the well-known toxicity of copper in many systems [8]. Novel in our findings are the following: First, the results indicate that transformed (3G) cells, express extremely fine and variegated cellular processes extending from the cells bodies. These processes take up large areas in the microenvironment compared to normal chondrocytes.

Second, over time, the transformed 3G cells demonstrated shedding of the extended fine processes, with release of material into the cell culture supernatant. When assayed for collagen content, it was found that this culture media had significantly elevated levels of collagen compared to normal culture media, indicating the potential transformation of 3G cells into functional producers of elevated extracellular matrix components.

Finally, it was found that 3G cells were stably transformed in the sense that when they were re-combined with normal chondrocytes, cellular boundaries and distinct morphologies for the two cells types remained. 3G cells continued in mixed cultures to take up more cell area and to express extended and variegated processes compared to smaller and simpler normal chondrocytes. This recombination of transformed 3G cells and normal chondrocytes may point to the potential for re-integration of 3G cells into normal tissues and for tissue engineering purposes.

We do not yet know the mechanism of transformation of chondrocytes by CuHARS treatment. Previously, others have used viral-mediated mechanisms to transform chondrocytes [9-11]. However, where available, these reports demonstrate no, or only modest morphological changes with increased proliferative rate compared to the normal parent cells [12]. No previous work to our knowledge has shown diminished proliferation, nor the extreme and variegated cellular process formation with membrane shedding, as we have shown here. One previously published work showed shedding and fragmentation of cell membranes similar to that shown here, but it was carried out in cancer cells adhered to collagen coated surfaces [13]. It is thus possible, that the indications of increased collagen production measured here, may be part of the mechanism for cell membrane shedding and fragmentation.

One anticipated possibility when we initiated these studies was that copper-containing CuHARS might be toxic to the cells, killing them, rather than upregulating them. Surprisingly, that was not observed, and thus one benefit of using CuHARS as a transforming agent may be the slow degradation rate of the material [4], and the lower toxicity compared to copper nanoparticles [14]. Copper salts have also been used previously to study effects on chondrocytes [6], but as completely soluble materials, the dynamic effects on cells would be hypothesized to be different than for the slowly degrading CuHARS used in the current work. Here we were able to track and quantify CuHARS materials changes associated with chondrocyte cultures over an 18-day period. Considering the utilization of copper by the collagen-stabilizing enzyme lysyl oxidase [5, 15] our results may be consistent with discovery and selection of transformed chondrocytes that generate and shed collagen that can be detected in cell culture supernatants (**Figure 2C**).

A limitation of this study is the low proliferation rate of transformed 3G cells compared to normal chondrocytes. Furthermore, to date, we have not been able to successfully cryopreserve transformed 3G cells but have passaged more than 20 flasks and as shown, have also now combined 3G cells with normal human chondrocytes. A challenge for cryopreservation with 3G cells may be that the extremely long and variegated processes of these cells are necessarily sheared by trypsin/EDTA treatment used to lift the cells off plating surfaces before concentrating for freezing/cryopreservation, which is a typical method for cryopreservation of cells [16]. We also used DMSO as the cryo-protective agent, but alternatives to this reagent are being explored for cell freezing [16]. In the current studies we also treated human dermal fibroblasts with CuHARS and found an attenuated morphology change compared to chondrocytes, but with retention of enhanced proliferation compared to 3G transformed cells (**Figure S4**). Furthermore, these treated fibroblasts were able to survive cryopreservation, which might also be consistent with their retained higher proliferation rate compared to 3G cells. Studies are underway to investigate methods for sustaining transformed 3G cells using alternative cryopreservation techniques along with growth factors and supplements, so that potential research questions of interest can be extended and shared across collaborative connections.

************************************

***********************************************

## Supplementary Material

